# Integrated Computational Biophysics approach for Drug Discovery against Nipah Virus

**DOI:** 10.1101/2023.10.23.563595

**Authors:** Georcki Ropón Palacios, Manuel Chenet Zuta, Jean Pierre Ramos Galarza, Edinson Gervacio Villarreal, Jhon Pérez Silva, Kewin Otazu, Ivonne Navarro del Aguila, Henry Delgado Wong, Frida Sosa Amay, Nike Dattani, Ihosvany Camps, Rajesh B. Patil, Abu Tayab Moin

## Abstract

The Nipah virus (NiV) poses a pressing global threat to public health due to its high mortality rate, multiple modes of transmission, and lack of effective treatments. NiV glycoprotein G (NiV-G) emerges as a promising target for NiV drug discovery due to its essential role in viral entry and membrane fusion. Therefore, in this study we applied an integrated computational and biophysics approach to identify potential inhibitors of NiV-G within a curated dataset of Peruvian phytochemicals. Our virtual screening results indicated that these compounds could represent a natural source of potential NiV-G inhibitors with ΔG values ranging from -8 to -11 kcal/mol. Among them, Procyanidin B2, B3, B7, and C1 exhibited the highest binding affinities and formed the most molecular interactions with NiV-G. Molecular dynamics simulations revealed the induced-fit mechanism of NiV-G pocket interaction with these procyanidins, primarily driven by its hydrophobic nature. Non-equilibrium free energy calculations were employed to determine binding affinities, highlighting Procyanidin B3 and B2 as the ligands with the most substantial interactions. Overall, this work underscores the potential of Peruvian phytochemicals, particularly procyanidins B2, B3, B7, and C1, as lead compounds for developing anti-NiV drugs through an integrated computational biophysics approach.

**Key points:**

1. **Nipah Virus (NiV) Threat:** NiV is a severe public health risk due to its high mortality rate, broad host range, multiple transmission modes, and lack of effective treatment. Outbreaks have occurred frequently in South and Southeast Asia, particularly in Bangladesh and India, leading to high fatality rates.
2. **Cross-Border Concerns:** NiV’s ability to transmit between humans and domestic animals raises concerns about its potential to cross regional borders and cause pandemics. It has been recognized as a high-priority pathogen by the World Health Organization.
3. **Lack of Treatment:** Currently, there are no approved specific antiviral treatments or vaccines for NiV. Patients receive supportive care and some drugs used for other viruses, despite their side effects.
4. **Targeting NiV Glycoprotein G:** The study focuses on NiV glycoprotein G (NiV-G) as a target for potential anti-Nipah drugs due to its crucial role in viral entry. This glycoprotein mediates viral attachment and entry into host cells.
5. **Computational Drug Discovery:** The research employs computational methods, including virtual screening and molecular dynamics simulations, to identify potential inhibitors of NiV-G from a dataset of Peruvian phytochemicals, particularly procyanidins B2, B3, B7, and C1. These compounds showed promising binding affinities, stable interactions, and favorable binding energies with NiV-G, making them potential lead compounds for drug development.

## 1. Introduction

Nipah virus (NiV) poses a dreadful risk to global public health due to its high mortality, wide host range, multiple modes of transmission, and lack of effective treatment. Since its first emergence in 1998, several outbreaks have occurred almost yearly in South and Southeast Asia countries, particularly in Bangladesh and India [1]. Despite being seasonal and sporadic, these outbreaks have resulted in numerous human deaths, with a high case fatality rate of over 73% [2, 3]. This infection can lead to severe respiratory illness, multiple organ failure, deadly encephalitis and seizures, ultimately resulting in coma [4]. The spread of NiV infection from Malaysia to other countries can be ascribed to the extensive geographic distribution of its natural reservoir, Pteropus fruit bats [5]. Infected bats can transmit NiV to humans by direct exposure to bat secretions on fruit and date palm sap or via a wide range of symptomatic intermediate host animals such as pigs, horses, dogs, and cats [6].

These modes of intraspecific (human-human) and interspecific (domestic animal-human) transmission have raised global concern about NiV’s potential to cross regional borders and cause subsequent pandemics [7]. As a result, it has been recognized as a high-priority pathogen by the World Health Organization R&D Blueprint to develop countermeasures for a fast and effective epidemic and pandemic response [8]. However, as NiV must be handled in BSL-4 containment laboratories, only a limited number of laboratories worldwide are capable of working with the wild-type virus [7].

Since there are no approved specific antiviral treatments or vaccinations for NiV, current patients only have access to supportive care and some drugs previously used against other paramyxoviruses (e.g., Ribavirin, Acyclovir). Although these drugs are not proven treatments for NiV, they are currently employed as a first-line strategy in emergencies for acute NiV encephalitis [9] despite their side effects, such as nausea, vomiting, and convulsions. Hence, exploring novel small-molecule chemical inhibitors or phytochemicals against NiV that hold potential therapeutic value is imperative while minimizing or eliminating side effects in humans.

In this sense, the NiV glycoprotein G emerged as a promising target for the identification of potential specific anti-Nipah drugs due to its crucial role in NiV viral entry. The negative-sense single-stranded RNA genome of NiV encodes six structural proteins, including G and F glycoproteins. These viral glycoproteins are essential for the recognition of host-cell-surface receptors ephrin-B2 (EFNB2) and ephrin-B3 (EFNB3), which mediate cellular attachment, fusion, and virus entry. NiV glycoprotein G (NiV-G) has a globular head domain formed of a six-bladed beta-sheet propeller connected via a flexible stalk domain to a transmembrane anchor [10]. The binding of G glycoprotein to its ephrin receptors leads to conformational changes in this glycoprotein, followed by subsequent conformational changes in F glycoprotein that facilitate the fusion of viral and cellular membranes [11,12]. Furthermore, NiV-G/F glycoprotein complexes expressed on infected cell surfaces also facilitate cell-to-cell fusion with neighboring non-infected cells, promoting the spread of infection [13].

Based on these observations, it was hypothesized that inhibitors targeting the active site of NiV-G responsible for host receptor interaction may disrupt viral entry into the cell. However, the considerable costs associated with establishing and maintaining an experimental high-throughput screening platform for identifying NiV-G inhibitors hinder their practicality for drug discovery. Additionally, the scarcity of BSL-4 laboratories worldwide further constrains the exploration of novel potential NiV inhibitors. Indeed, computational approaches now enable the virtual screening of millions of compounds in less time, thereby reducing the initial costs of hit identification, filtering the pool of drug candidates to a fraction with the most promising hits, and improving chances of discovering viable drug candidates during experimental validation [14].

Pursuing this aim, the present study conducted a computational-guided drug discovery based on virtual screening and molecular dynamics (MD) simulations to identify novel potential inhibitors of the glycoprotein G of Nipah virus (NiV-G) within a curated dataset of Peruvian phytocompounds.

## 2. Methods

### 2.1. Receptor and ligand preparation

The three-dimensional structure of the NiV-G protein (receptor) with a resolution of less than 2.3 Å, was obtained from Protein Data Bank (ID: 3D11). The missing atoms of the structure were added by PDBReader [15] from the CHARMM-GUI web tool [16], and the protein was protonated at Ph 7.0 with the PDB2PQR tool [17]. The ligands of our in-house custom database of Peruvian phytochemical compounds were prepared with GypsumDL [18]. The smiles of these ligands were converted to their 3D structure in PDB format with a pH range of 7.4-8.4 and 5 conformomers per each ligand. The Durrant filter implemented in GypsumDL was applied to the ligands during the conversion. Finally, the PDB of the receptor and ligands were converted to PDBQT format using AutoDockTools [19].

### 2.2. Virtual screening and molecular docking

Active site determination was based in our previous work [20], where the following residues were identified as key for interaction with potential ligands: Asp219, Arg236, Pro220, Cys240, Pro276, Asn277, Val279, Tyr280, His281, Cys282, Tyr351, Gly352, Pro353, Phe458, Gly489, Gln490, Gly506, Val507, Tyr508 and Lys560. For docking we used a grid of size 22.13, 35.88, and 20.96, centered at 22.13, 35.88 and 20.96 (in the x, y, and z axis respectively) through an in-house Tcl script based on VMD [21] and an exhaustive search equal to 20 using Autodock-Vina software [22]. Once the active site and search box were obtained, a virtual screening was performed with the previously prepared library of 1415 conformers. The top 10 conformers were filtered based on its interaction profile with the receptor through PLIP analysis [23] and one conformer per ligand was selected for further characterization with MD simulations.

### 2.3. Biomolecular system setup

We prepared five systems, one apo and four in complex with selected procyanidins, using QwikMD [24]. The systems were solvated into a cubic box of 15 Å of padding with the TIP3 [25] water model, in the presence of 150 mM of NaCl to emulate the physiological conditions of the biological environment. The Ligands were parameterized with the CGenFF tool [26, 27]. All simulations were run with NAMD3 [28] and the CHARMM36 FF [29, 30].

### 2.4. Molecular dynamics simulation study

The constructed systems were minimized in the presence of 5 kcal mol^-1^ Å^2^ of harmonic restraint on heavy atoms to the ligand and backbone to the protein during 3000 steps. The system was gradually heated from 60 K to 310 K in NpT ensemble with a pressure of 1 atm, keeping the harmonic restraint during 5 ns, followed by equilibration into NpT ensemble with 310 K and 1 atm of pressure with the same restraint during 10 ns. Finally, the unrestrained production phase MD simulations of 300 ns were performed, and trajectories were collected at each 10 ps. The Lavengin thermostat and the Nosé-Hoover Langevin piston [31, 32] were used to control temperature and pressure in the equilibration and production steps. Rigid hydrogen atoms were kept with the SETTLE algorithm [33] in water, while the solute atoms were kept with the RATTLE/SHAKE algorithm [34]. A distance cut-off of 12 Å was applied to short-range, nonbonded interactions, and 10 Å for the smothering function. Long-range electrostatics interactions were treated using the particle-mesh Ewald [35] method. The r-RESPA multiple time step scheme [28] was used in all cases with 2 fs of time integration steps.

### 2.5. Non-equilibrium free energy calculation

For this point, the system was prepared and equilibrated as is described above, with QwikMD [24], except a 30 Å padding extra into the Z axis was added to enable pulling. For pulling a potential bias of the form 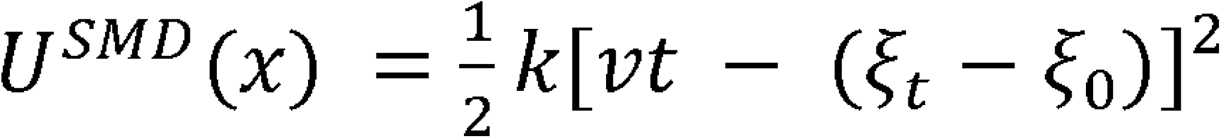 was added for the original potential *H*^±^ = *V*^*original*^ + *U*^*SMD*^ through the colvars module [36], getting a modified potential (*H*^±^). For this, the center of mass of the ligand, was pulling with a speed v of 10 Å ns^-1^ and of 100 kcal mol^-1^ Å^-2^ during 3 ns, enough to reach the unbound state. During the pulling the C_a_ atoms from protein were restrained with a force of 5 kcal mol^-1^ Å^-2^. A total of 7 independent replicates were performed (total 21 ns). The work was computed for numerical integration using 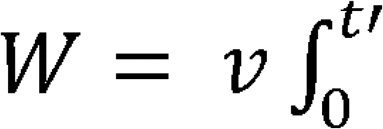*dt*′*F*(*t*′). From these simulations, the potential of mean force (*PMF* (*ξ*), *Φ* (*ξ*)) was reconstructed through Jarnzynski identity [37] as show in the following equations:

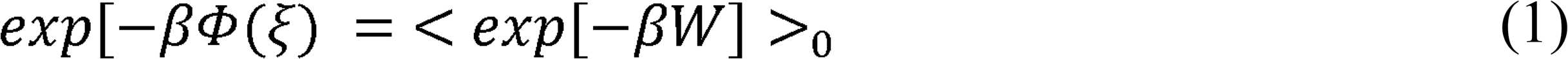

If rearranging the equation it shield the *Φ* (*ξ*):

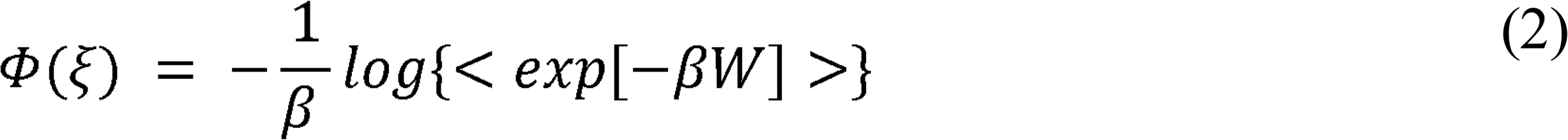

And expanding this equation for the 2nd order cumulant:

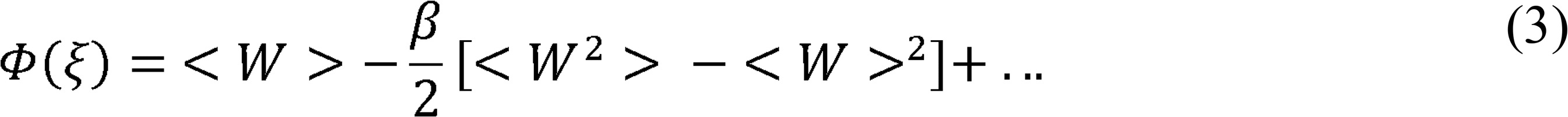

Where *Φ* (*ξ*) is the potential of mean force, i.e the free energy profile over the collective variable (*ξ*) *W* is the work done along *ξ*, and *β* is the inverse of thermal energy (1/*k* _B_T), where *k*_B_ is Boltzmann’s constant.

On the other hand, to calculate the 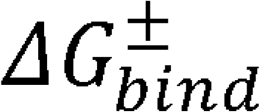 standard, we used the following expression:

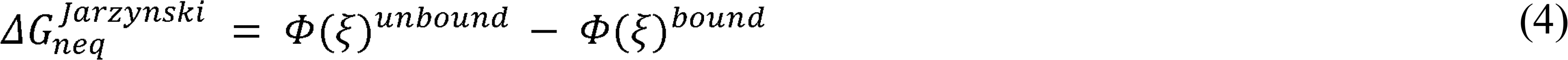

Then

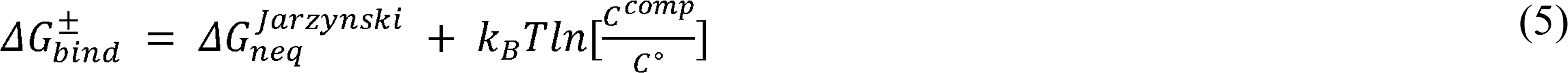

Where 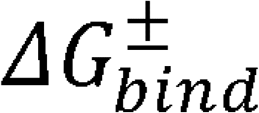 in kcal mol^-1^, in kcal mol^-1^ K^-1^, *C*^*comp*^ 1 ligand 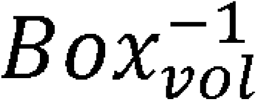 is the concentration used in the simulation in Å^3^ and *C*° 1/1661 Å^3^ is the standard concentration yielding 1 ligand x 1661 Å^-3^ . These last two terms were added to pass from the computational system to the standard system. Here *Φ* (*ξ*) ^*bound*^ and *Φ* (*ξ*) ^*unbound*^ correspond to *Φ* (*ξ*) the associated and dissociated thermodynamics state (i.e macrostate) respectively.

### 2.6. Molecular data analysis

All analyzes were performed through in-house Tcl/Python scripts based on VMD 1.9.4 [21] and MD analysis 2.0 [38] respectively. The 3D representations were rendered with VMD 1.9.4. The plots were generated with in-house R scripts with the ggplot2 [39] library. Interaction throughout the simulations were computed with an in-house R implementation of MD interaction plot tool (https://github.com/tavolivos/Molecular-dynamics-Interaction-plot).

## 3. Results and Discussion

### 3.1. Determination of NiV-G binding pockets

The NiV-G monomeric structure used in this study is represented in **Figure** 1 with its binding pocket highlighted in red. This glycoprotein presents a β-propeller fold with six blades formed by four-stranded antiparallel β-sheet surrounding a central cavity [40]. NiV-G binding pocket is located at the center of the top face of the β-propeller structure [40], and key residues used for the virtual screening are displayed in **Figure** 1.

**Figure 1.**
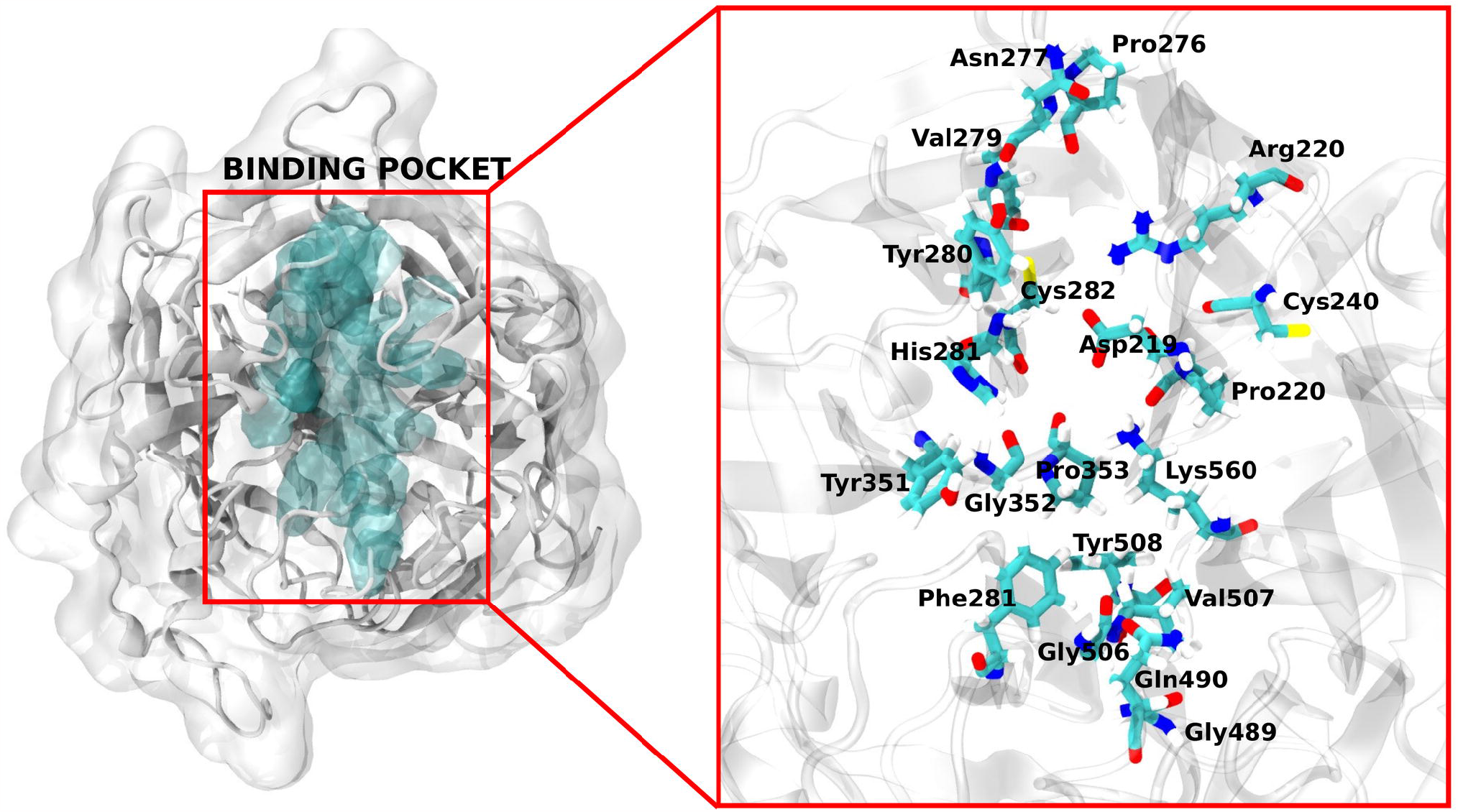
NiV-G monomer structure representation. Left: NiV-G structure with its pocket highlighted in skyblue. Right: Top view of NiV-G Binding pocket with key residues displayed in sticks.

### 3.2. Virtual Screening and molecular docking study

Virtual screening results highlight the potential of Peruvian phytochemicals for anti-Nipah drug discovery. Most conformers screened against NiV-G exhibited binding energies lower than -6 kcal/mol (**Figure 2A**). Around 40% of the conformers displayed a favorable binding energy exceeding -8 kcal. These binding affinities suggest that these diverse phytochemicals share structural patterns that could be useful as scaffolds for drug design in future studies.

**Figure 2.**
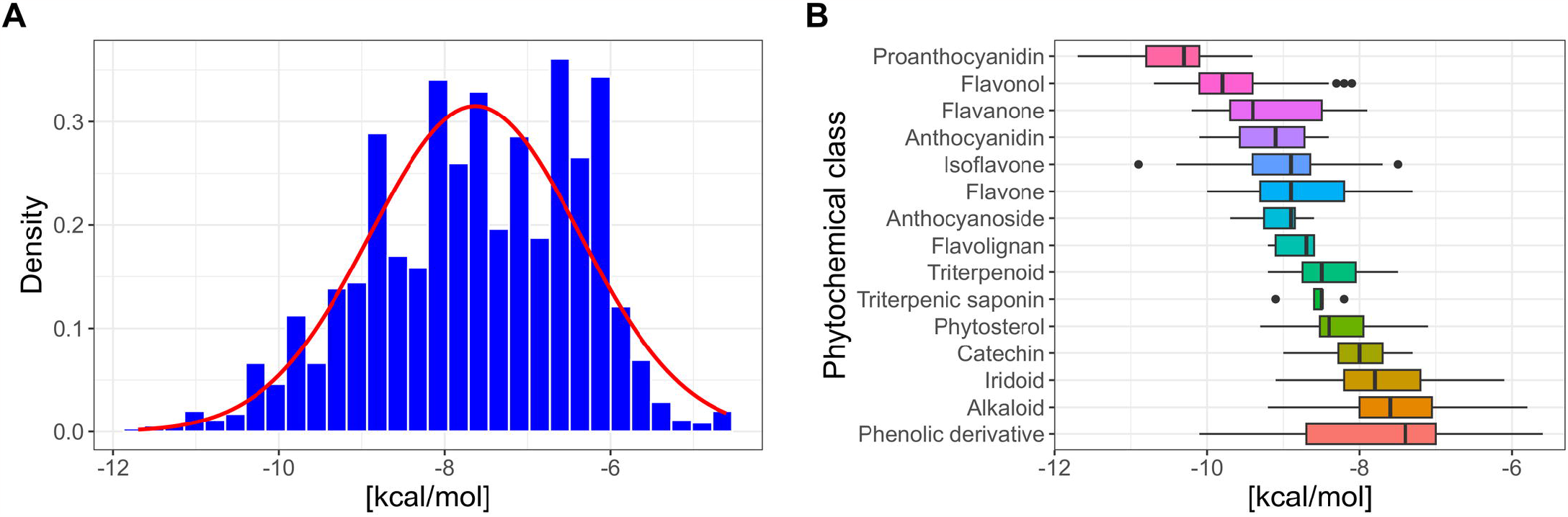
Virtual screening of peruvian phytochemicals against NiV-G. (**A)** Binding energy probability distribution of conformers screened. (**B)** Boxplot of grouped conformers by phytochemical class. Proanthocyanidins conformers exhibit the highest binding energies.

Procyanidins stand out as the phytochemical class with the highest binding forces with NiV-G followed by several flavonoids and derived compounds. These plant secondary metabolites have well-documented anti-inflammatory [41], anti-infectious [42], antimicrobial [43, 44], and antiviral [45] properties. Computational investigations have further identified flavonoids as promising sources of potential inhibitors against various viral targets, such as the main protease of SARS-CoV-2 [46, 47], Dengue Non-Structural protein 1 (NS1) [48], and HIV-1 protease [49], among others.

Procyanidins B2, B3, B7, and C1 exhibited the highest binding energy values (around -11 kcal/mol) and the best interaction networks with NiV-G (**Figure** 3, **Table** 1). Notably, these compounds contain flavan-3-ol monomers as basic units in their structure, which are composed exclusively of (+) catechin and (–) epicatechin [50]. Therefore, they are characterized by the presence of multiple aromatic rings with hydroxyl groups that allow interactions with proteins through hydrogen bridges [50]. These chemical characteristics make procyanidins an attractive target for drug design approaches such as ligand-based pharmacophore modeling [51].

**Figure 3.**
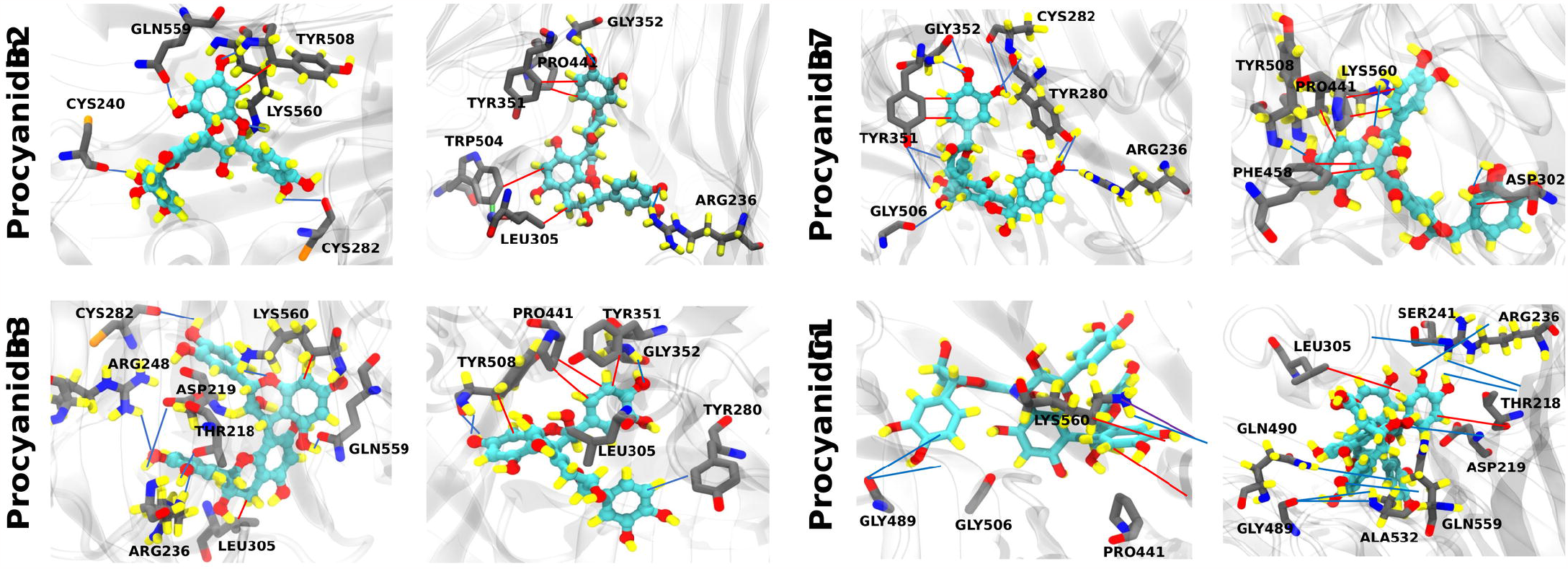
Representation of molecular interactions between selected procyanidins and NiV-G binding pocket. The ligands (skyblue) and the binding pocket of NiV-G (dark gray) were drawn using licorice representation in VMD. Interactions are represented as lines of different colors indicating the presence of hydrogen bonds (blue), hydrophobic interactions (red), pi stacking (purple). Atoms follow a color code indicating the presence of nitrogen (blue), oxygen (red), sulfur (orange) and hydrogen (yellow).

**Table 1.**
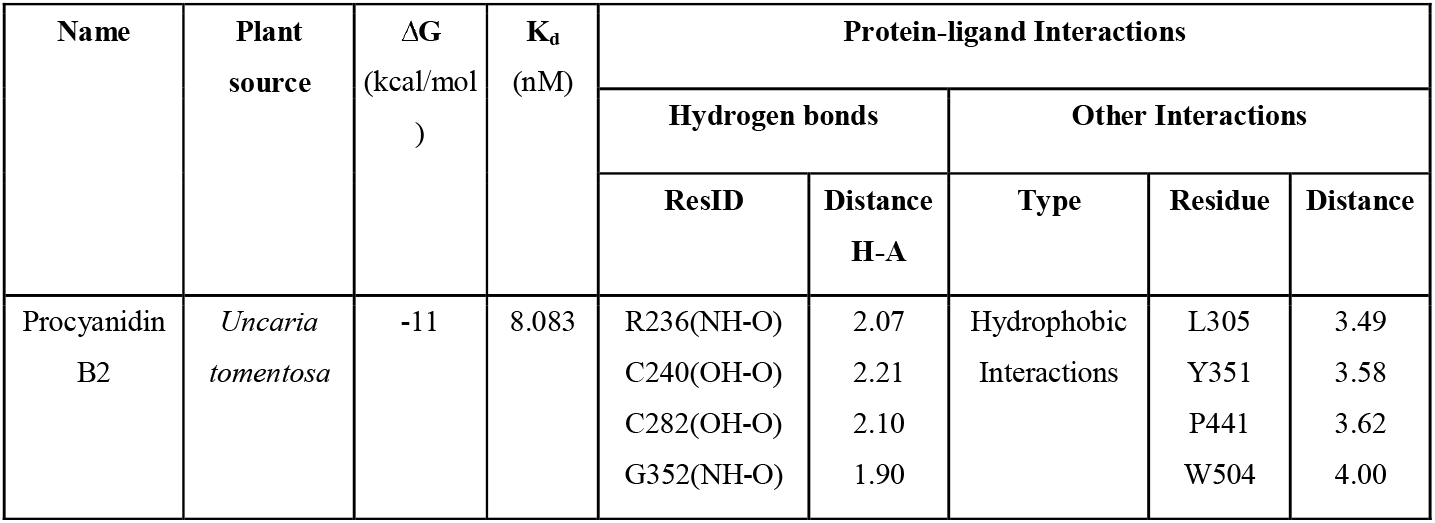
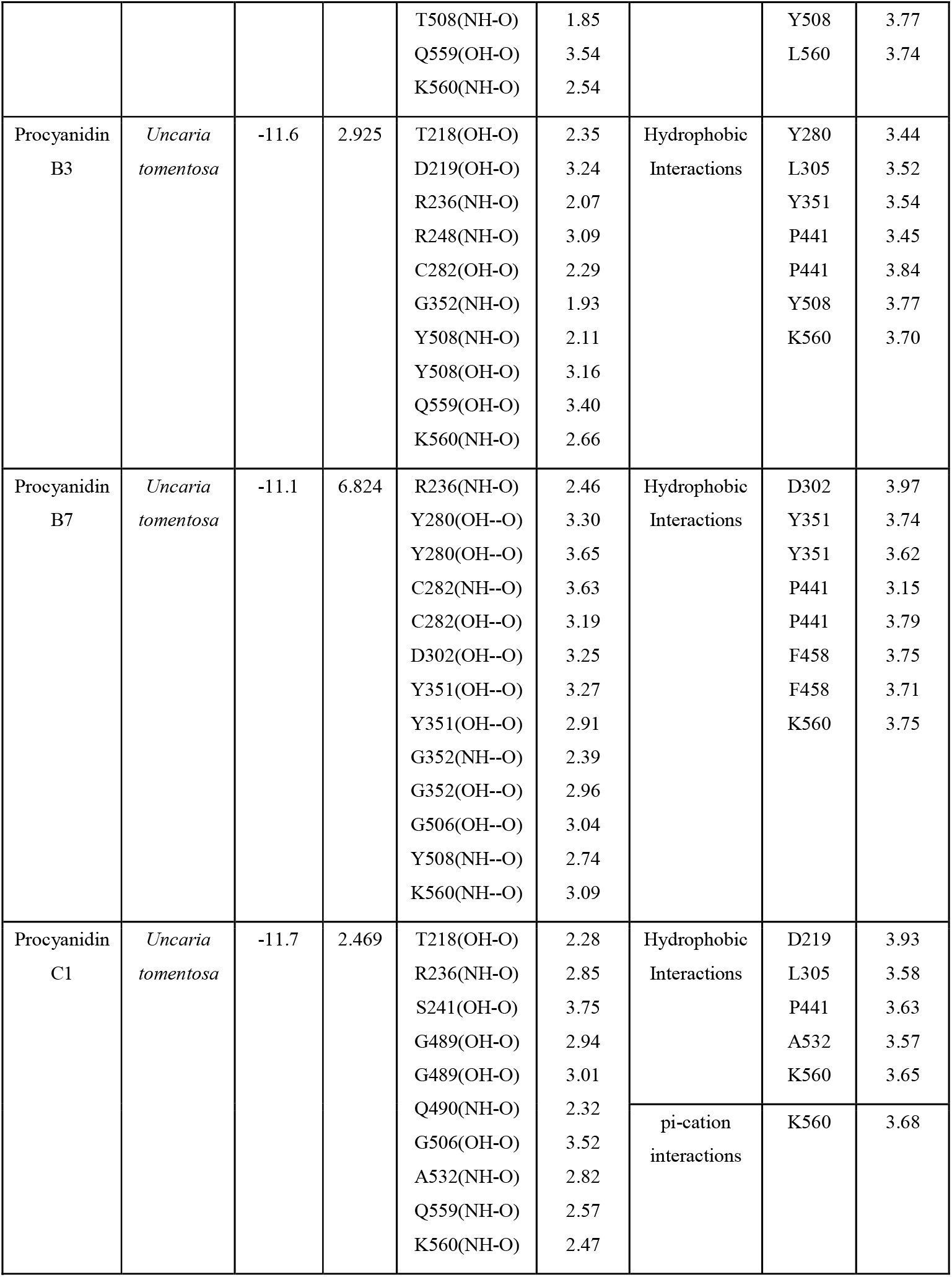
Top ranked phytochemicals screened against Nipah virus G glycoprotein (NiV-G).

Although Procyanidin B7 doesn’t exhibit the highest binding energy (-11.1 kcal/mol) among the four chosen ligands, it stands out for having the most extensive array of molecular interactions with NiV-G (13 hydrogen bonds and 8 hydrophobic interactions).

In contrast, Procyanidin C1 displayed the most favorable binding energy (-11.7 kcal/mol) compared to all other ligands, but it only engaged in 10 hydrogen bonds and 5 hydrophobic interactions with residues D219, L305, P441, A532, and K560, along with a single π-cation interaction with K560. Procyanidin B3 emerged as the second ligand with the highest binding affinity at -11.6 kcal/mol, forming 10 hydrogen bonds and 7 hydrophobic interactions. Finally, Procyanidin B2 displayed a favorable energy value of -11.1 kcal/mol, establishing 7 hydrogen bonds and 7 hydrophobic interactions.

These interaction profiles underscore the high affinity of selected procyanidins for NiV-G pocket. There is a shared binding pattern among the selected procyanidins within the NiV-G pocket. Notably, specific residues such as R236, Y280, C282, G506, and Q559 consistently engage in hydrogen bond formation with the ligands. On the other hand, residues like L305, Y351, P441, and F458 are recurrent across all ligands hydrophobic interactions, suggesting their key role in the binding mechanism between NiV-G and procyanidins.

In fact, NiV-G binding pocket has been described as a hydrophobic pocket [40, 52, 53] that recognizes and interacts with Ephrin B2/B3 accommodating hydrophobic residues of host receptors through an induced-fit mechanism. Specifically, T218, L305, G489, Q490, and A532 residues were identified as critical for NiV-G binding to Ephrin B3 external loops [40]. Additionally, Q559 residue participates along with G579, Y581, and I588 on Ephrin B2 binding pocket of NiV [54]. Therefore, further studies should be addressed to probe that these procyanidins could also work as competitive inhibitors of NiV-G binding with Ephrin B2 and B3 host receptors.

Therefore, for further understanding of procyanidins impact on NiV-G structural features such as structural stability, conformational changes, mobility fluctuations, and collective motions, we processed our MD trajectories with root-mean-square deviations (RMSD) (**Figure** 4), root-mean-square fluctuations (RMSF) (**Figure** 5) and principal component analysis (PCA) (**Figure** 6) analysis. On the other hand, we employed 2D-FEL using RMSD and radius of gyration (Rg) (**Figure** 7), PLIP occurrence (**Figure** 8), and visual representations (**Figure** 9) of trajectories to evaluate the preferred conformational states of procyanidins and their binding mechanisms with the NiV-G pocket.

**Figure 4.**
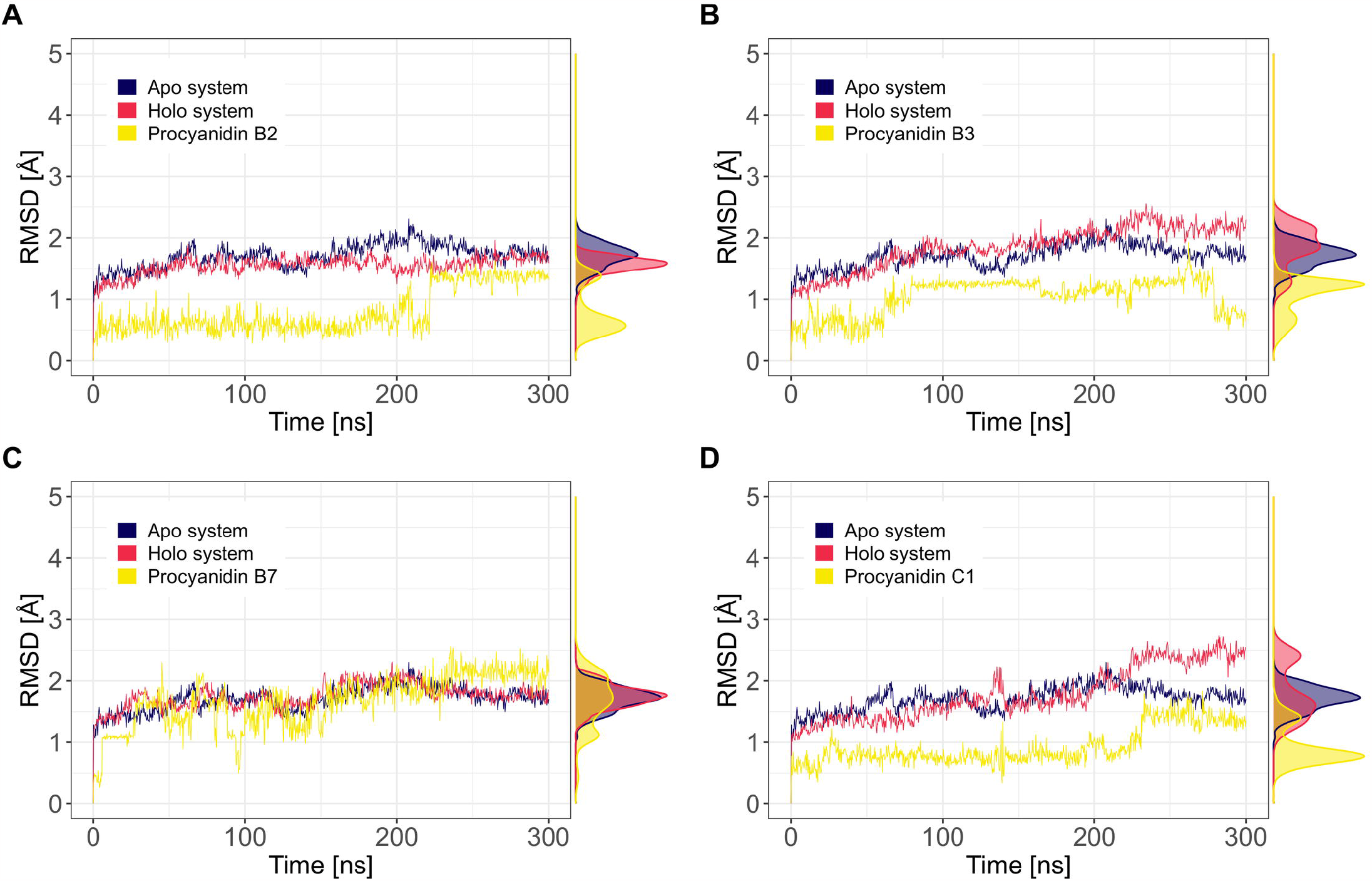
RMSD of NiV-G complexes with Procyanidin B2 **(A)**, B3 **(B)**, B7 **(C)**, and C1 **(D)**. Lines represent RMSD values of NiV-G simulation (dark blue), complexes simulations (red) and ligands from complexes simulations (yellow).

**Figure 5.**
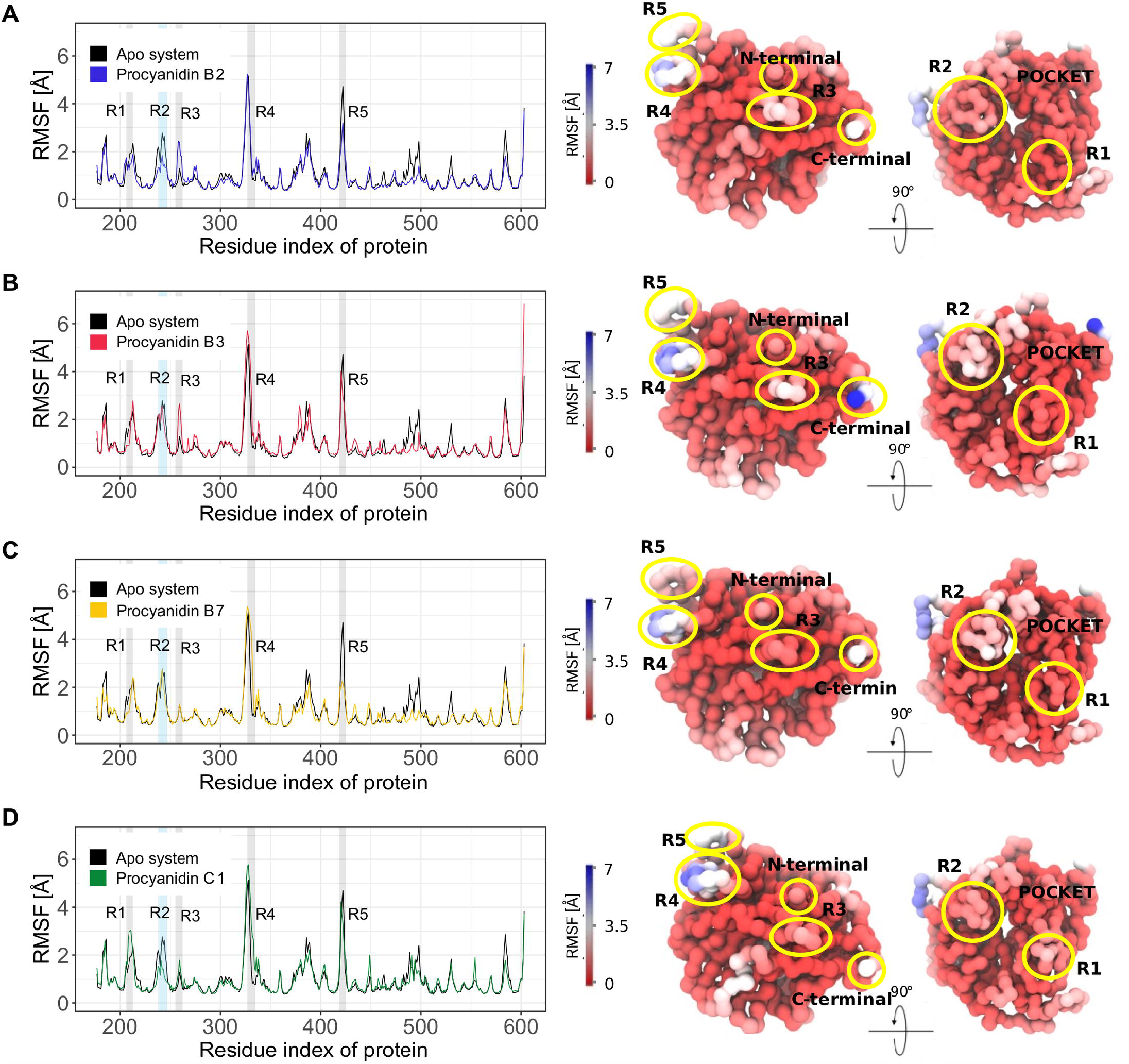
RMSF of NIV-G residues of NiV-G in complex with Procyanidin B2 **(A)**, B3 **(B)**, B7 **(C)**, and C1 **(D)**. Left: RMSF values of apo and holo systems for each procyanidin. Divergent regions and pocket residues are highlighted with gray and sky blue boxes respectively. Right: 3D representation of RMSF values as β factors, where red indicates the most stable regions and blue the most fluctuating ones. Divergent regions are indicated with yellow circles.

**Figure 6.**
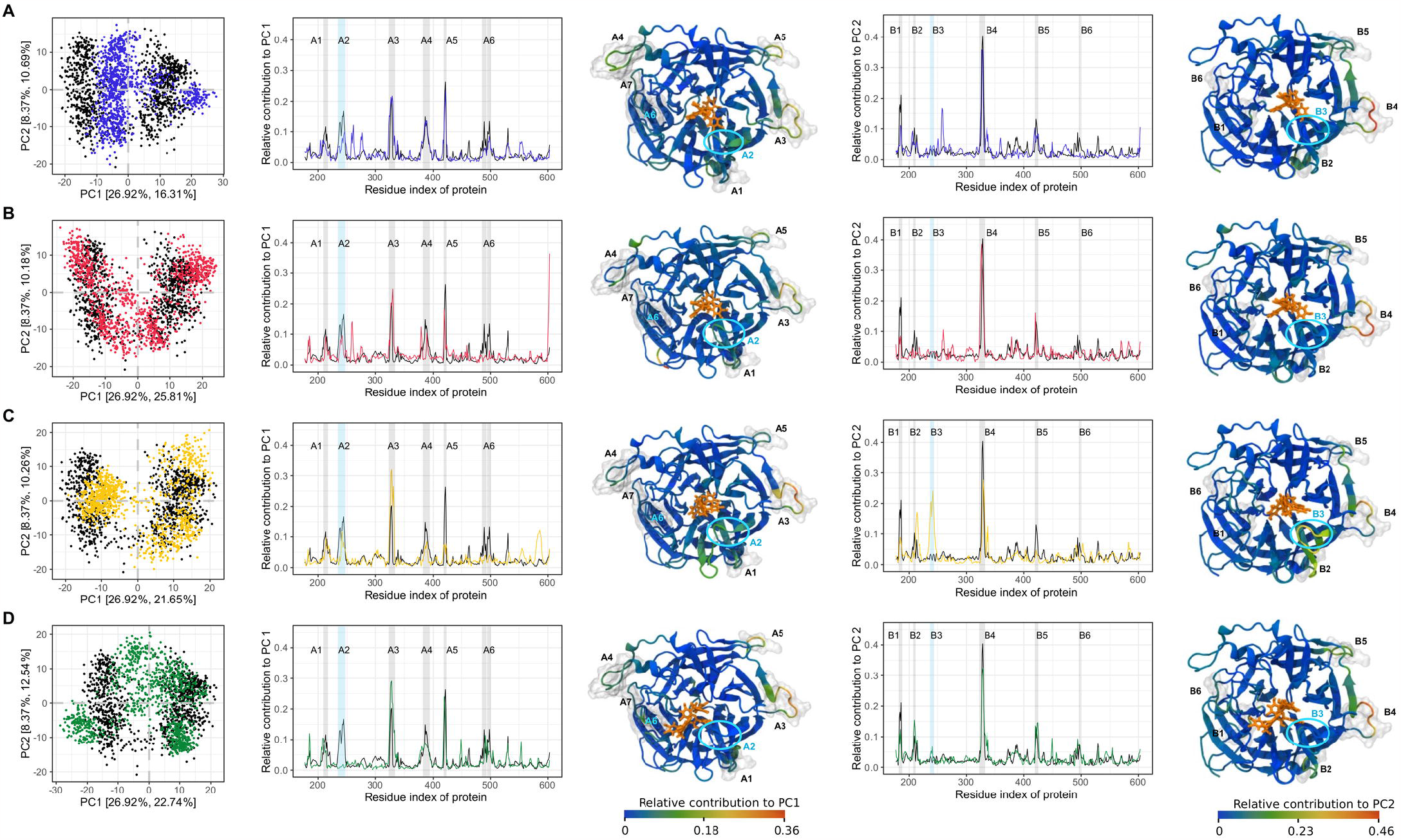
PCA of NiV-G complexes with procyanidins B2 (A), B3 (B), B7 (C), and C1 (D). From left to right: I) the conformer plots of NiV-G structures defined by PC1 and PC2. II) Relative contributions of residues to PC1. III) Representation of the collective motion defined by PC1. IV) Relative contributions of residues to PC2. V) Representation of the collective motion defined by PC2.

**Figure 7.**
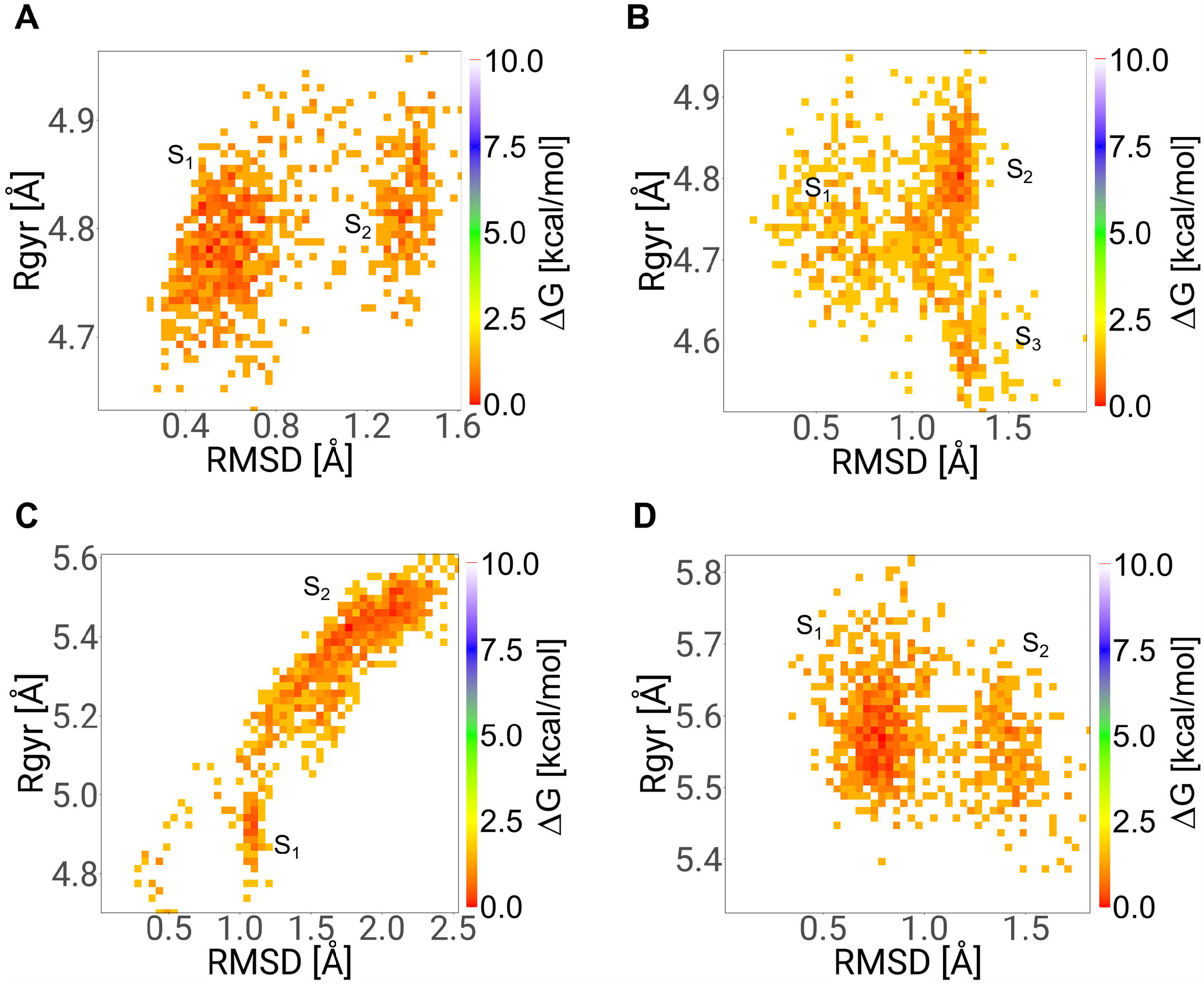
Conformational states adopted by Procyanidins upon binding with NiV-G pocket. 2D Free Energy Landscape plots of procyanidin B2 **(A)**, B3 **(B)**, B7 **(C)**, and C1 **(D)** based on RMSD and Rg values. Favorable conformational states of ligands are denominated as S1, S2 and S3.

**Figure 8.**
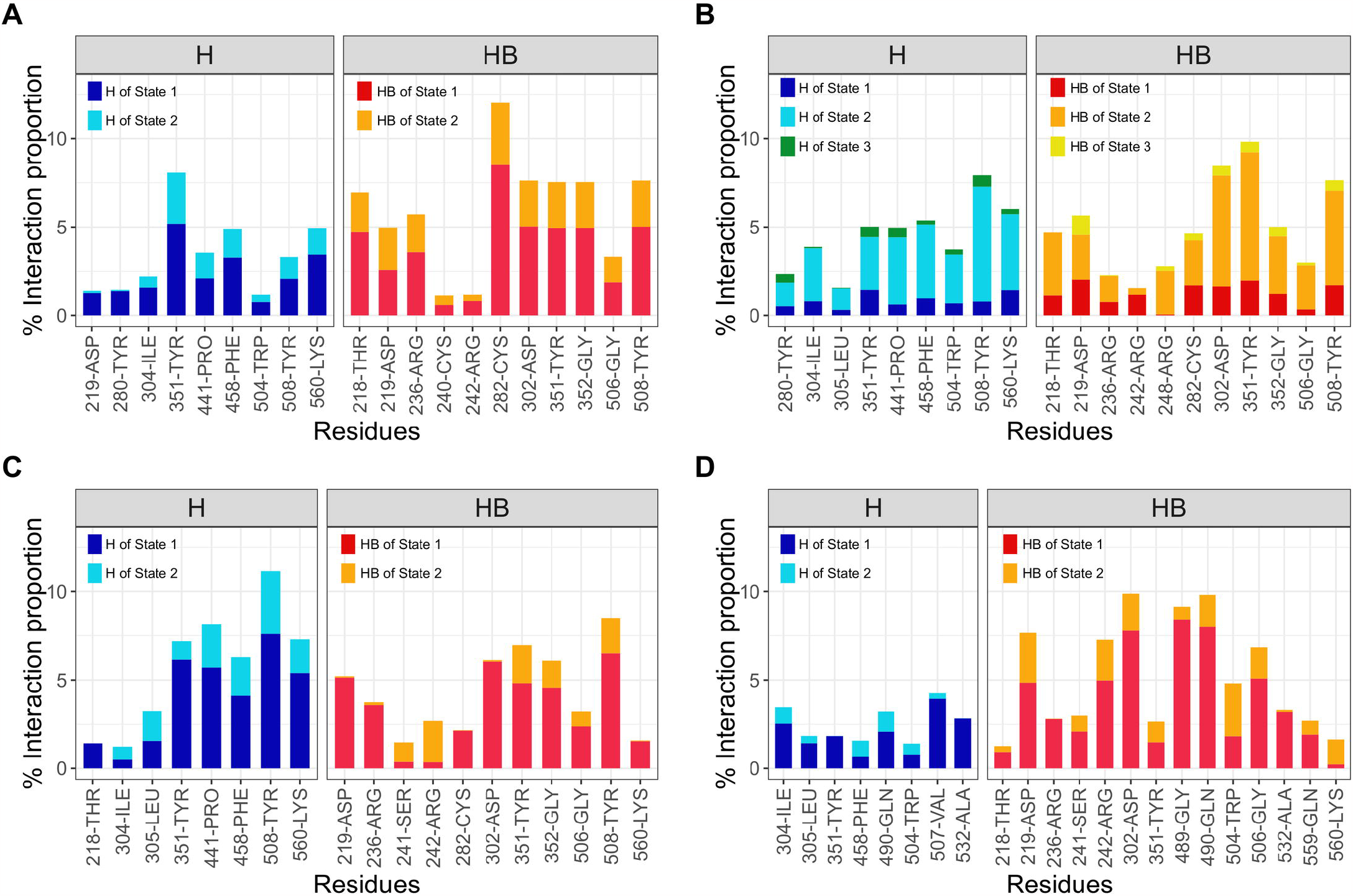
Molecular interactions of procyanidins and NiV-G. **(A)** Procyanidin B2 interactions with NiV-G present in its state 1 (0-220 ns), and 2 (220-300 ns). **(B)** Procyanidin B3 interactions with NiV-G present in its state 1 (0-90 ns), state 2 (90-280), and state 3 (280-300 ns). **(C)** Procyanidin B7 interactions with NiV-G present in its state 1 (0-230 ns) and state 2 (230-300 ns). **(D)** Procyanidin C1 interactions with NiV-G present in its state 1 (0-230 ns) and state 2 (230-300 ns). Interaction proportions were calculated based on the preferred conformational states previously identified with FEL. Time ranges for each conformational state of ligands were estimated based on its RMSD values. The interactions are clustered by type and shown in bar plots including hydrophobic (H) and hydrogen bonds (HB) with different colors.

**Figure 9.**
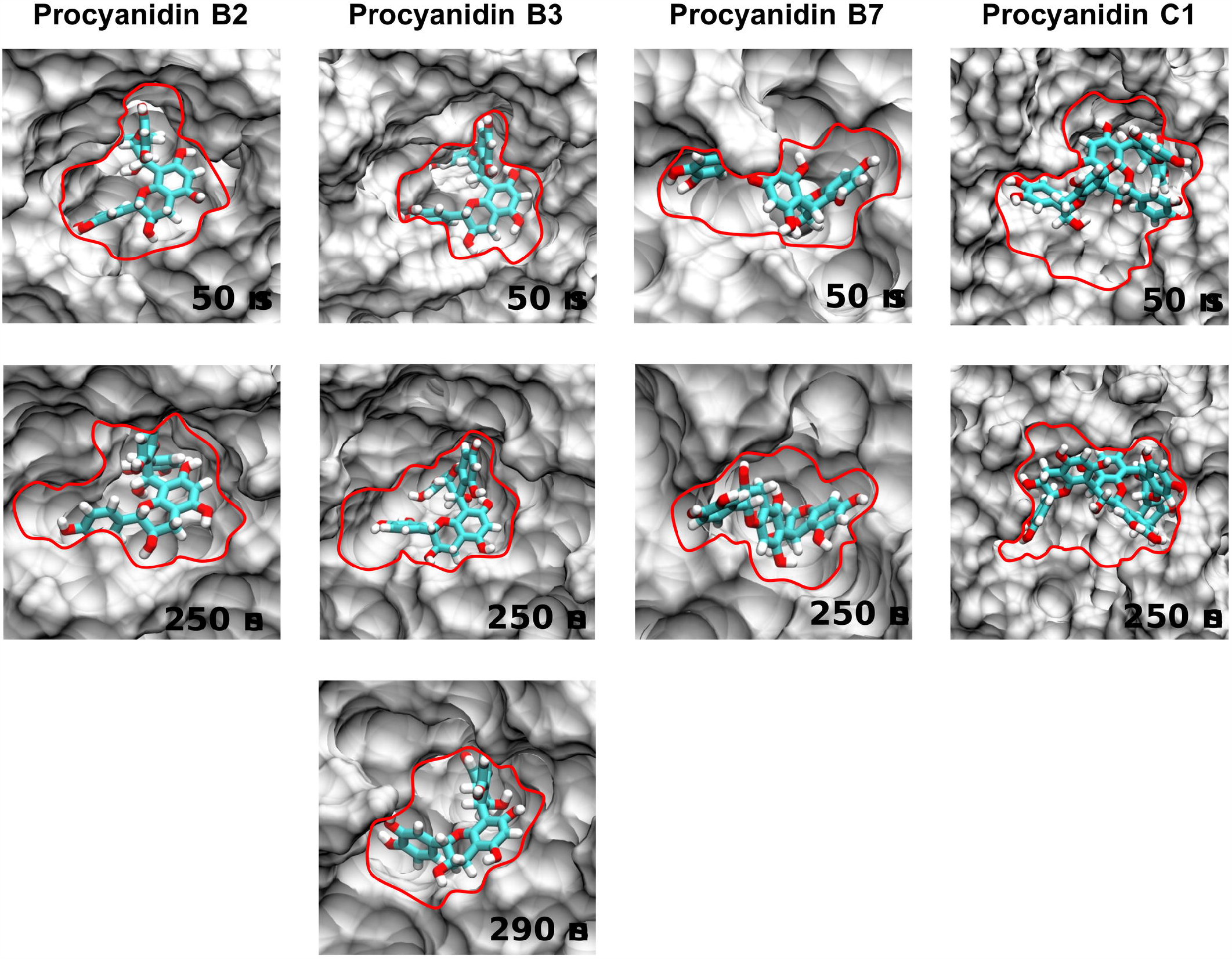
Temporal evolution of NiV-G pocket induced-fit interaction with procyanidins B2, B3, B7 and C1. The image shows the ligand interaction area in the NiV-G pocket and the conformational states of the ligand at different times of the MD simulation.

### 3.3. Molecular dynamics simulation study

#### 3.3.1. Analysis of root-mean-square deviations

The stability and structural changes of NiV-G-phytochemical complexes were evaluated through RMSD. The RMSD is the average displacement of the atoms at an instant of the simulation relative to a reference structure, usually the first frame of the simulation or the crystallographic structure [55]. This method is useful to understand time-dependent motions of the structure, such as its stability along the time of simulation or the conformational perturbation induced by a ligand.

According to our RMSD results (**Figure** 4), procyanidins do not significantly impact the complex stability along the 300 ns of simulation. NiV-G apo system achieved an average RMSD value of 1.71 Å while holo systems exhibited a slight variations with average RMSD values of 1.55, 1.85, 1.74, and 1.79 Å for procyanidins B2, B3, B7 and C1 respectively. Overall, NiV-G and procyanidins complexes remain stable during the 300 ns of simulation, with some minor conformational changes experimented upon ligand binding.

Among the ligands, procyanidins B2 and B7 present the more stable interaction with NiV-G (**Figure** 4A, C). On the other hand, Procyanidin B3 and C1 induce a slight perturbation on the complex stability (**Figure** 4B, D) with an increment of almost 0.5 and 1Å respectively. Further analysis (**Figure** 7, 8 and 9) suggested that these conformational changes are triggered by ligands interaction, as part of the induced-fit mechanism of NiV-G binding pocket.

Compared to our previous study [20], selected procyanidins exhibited more stable interactions with NiV-G structure than drugs MMV020537, MMV019838, and MMV688888 of the Pathogen Box. Similarly, other studies that applied MD simulations to assess NiV-G stability in complex with ligands, have reported RMSD values of 1.5 up to 2 Å for 100 ns simulation with 5 potential inhibitors [56], and 2 up to 3 Å with 2 potential peptide-based inhibitors [57]. Altogether, our RMSD results confirmed that proposed potential inhibitors do not significantly affect the global structural stability of NiV-G upon binding.

#### 3.3.2. Analysis of root-mean-square fluctuations

For further assessment of procyanidin’s impact on structural dynamics of NiV-G, we conducted the RMSF analyses of protein backbone α-carbons. RMSF is a measure of the displacement of a particular atom, or group of atoms, relative to the reference structure, averaged over the number of atoms [55]. Accordingly, when an area of the structure presents high RMSF values, it frequently diverges from the average, indicating high mobility.

Our RMSF results (**Figure** 5) identified five divergent regions, namely R1, R2, R3, R4 and R5, in NiV-G-procyanidins complexes. RMSF values of these regions fluctuated for almost 1 Å, suggesting potential conformational adaptations induced by the ligands. Most residues of NiV-G remain rigid during the simulation, corroborating its overall conformational stability.

On the bottom part of NiV-G structure, we identified other two divergent but also highly flexible regions, R4 (residues 329 to 333) and R5 (residues 420 to 423). These regions exhibited RMSF values of almost 5.5 Å, with the most increment upon procyanidins B3 and C1 binding.

Particularly, the NiV-G binding pocket cavity exhibited varying degrees of flexibility upon ligand binding. Two divergent regions, R1 (residues 208 to 211), and R3 (residues 257 to 260), were located closely to this structure while R2 (residues 240 to 245) was identified within this pocket, comprising one of its key residues, C240 (**Figure** 1). Specifically, R1 and R2 exhibited a flexibility exchange as part of the local changes introduced by the selected procyanidins.

In the presence of procyanidins C1 and B2, R1 region exhibits lower mobility, while R2 adopts higher flexibility and vice versa, as observed with procyanidins B3 and B7, while R1 exhibits higher flexibility, R2 becomes more rigid. This mechanical tradeoff between R1 and R2, suggests that while some residues of NiV-G pocket accommodate to fit the ligand, others become more rigid to stabilize the interaction.

Overall, our results are in accordance with RMSF analysis of other NiV-G systems with potential ligands [20, 56, and 57]. Multiple of the divergent regions identified in this work are recognized and mentioned in those studies with some variations on residues selection range. In sum, selected procyanidins introduce multiple local changes in the dynamics of NiV-G structure. Specifically, we highlight the role of R1, R2, and R3 on the mechanical basis of NiV-G pocket induced-fit conformational rearrange, for further studies that would like to design allosteric inhibitors of NiV-G pocket.

#### 3.3.3. Principal component analysis

As we observed these changes in the mechanical properties of specific regions of NiV-G, we conducted a PCA to explore the collective motions of NiV-G upon procyanidin binding and assess the significance of the previously identified regions in its pocket adaptation process. PCA is a multivariate statistical technique applied to systematically reduce the number of dimensions needed to describe protein dynamics through a decomposition process that filters observed motions from the largest to smallest spatial scales [58]. Through this technique, we can determine the essential space (ES), a subset of new coordinates obtained from atomic Cartesian coordinates of trajectories, able to describe the majority of fluctuations of the systems, and with this ES, the essential dynamics governing these motions.

After examining the PCA results (**Figure** 6) across the four NiV-G complexes with procyanidins B2, B3, B7, and C1, we identified multiple key regions exhibiting a divergent behavior on the principal component 1 (PC1) and the principal component 2 (PC2) upon ligand binding. These divergent regions exhibited a fluctuation of 10 % or more on its relative contribution to PCs variance. As the first PCs represent the most important collective motions of NiV-G, these divergent regions are key to understanding the nature of NiV-G pocket interaction with procyanidins.

In the case of PC1, we identified seven key regions, namely A1, A2, A3, A4, A5, A6 and A7, and in PC2, we identified another six key regions, namely B1, B2, B3, B4, B5, and B6. These regions not only contributed most to PCs variance but also exhibited a dynamic behavior upon procyanidins binding. Among these regions, A2 (residues 235 to 246), and B3 (residues 239 to 243) were located within NiV-G binding pocket, comprising its key residue C240. This subregion of NiV-G pocket was previously described as key along with R1 (**Figure** 5) for the mechanical plasticity evidenced by the RMSF analysis. Although not so evident with A1 region (residues 210 to 216), B2 region (residues 208 to 210) shed light on this process as seen on PC2 plots. When A2 and B3 exhibit its highest contribution to collective motion of procyanidins and NiV-G binding, B2 also exhibit an increment as seen in **Figure** 6C. Therefore, these results altogether suggest that the subregions of NiV-G conformed by residues 208 to 216 and 235 to 246 are key for NiV-G pocket plasticity and adaptability to procyanidins.

Notably, in each complex, NiV-G exhibited a distinct loop (residues 323 to 332) characterized by high flexibility and a significant contribution to the collective dynamics of PC1. Our observations suggest that this region, identified as A3 in PC1, B4 in PC2, and denoted as R4 in the RMSF analysis (**Figure** 5), plays a pivotal role in the conformational rearrangement experienced by the NiV-G pocket upon binding with the ligand. Due to its proximity to the A2 and B3 regions within the NiV-G pocket, this loop could function as an interdomain flexibility hotspot, facilitating dynamic movements of adjacent residues to enhance their interactions with procyanidins.

Consequently, these changes in the collective motions of specific regions of NiV-G suggest a conserved mechanism of interaction with the NiV-G pocket among procyanidins. As a result, future studies in rational drug design should take into consideration the use of procyanidins or their derivatives to develop effective therapeutics against NiV infection.

#### 3.3.4. Analysis of free energy landscape

To gain deeper insights into the interaction of NiV-G with the ligands, we explored the free energy landscape (FEL) of each ligand, using RMSD and Rg as key structural descriptors.

Our findings revealed that each selected procyanidin ligand had at least one highly favorable conformational state. Procyanidins B2 and C1 displayed similar behavior, each showing two different conformational states. Notably, these procyanidins maintained high stability (with RMSD values ranging from 0.4 to 1 Å) for the majority of the simulation (as shown in **Figure** 4A and D) when they adopted the first conformational state. As they transitioned to the second state, a more dynamic behavior was observed.

In contrast, procyanidin B3 presented a more complex binding profile with three distinct states, one of which displayed a remarkable ΔG value of 0. This pattern is further confirmed by the RMSD plot of procyanidin B3, where we identified the first conformational state (S1) from 0 to 60 ns, the second and most favorable state (S2) from 60 to 230 ns, and the last state (S3) from 230 ns to the end of the simulation.

Lastly, Procyanidin B7 presented two distinctive favorable states (S1 and S2). The first state represents the transitional state adopted by this ligand, while the second state represents the dynamic conformational states adopted by procyanidin B7 within the pocket.

#### 3.3.5. Analysis of molecular interaction of NiV-G with procyanidins

For further comprehension of procyanidin’s mechanism of interaction with the NiV-G pocket, we got the proportion of molecular interactions established between procyanidins and NiV-G during MD simulations (**Figure** 8) with an in-house script to run PLIP over each frame. In addition to that, we represented the preferred conformational states of ligands (**Figure** 9) selecting distinctive time points based on RMSD values of FEL. Each of these representations revealed distinct conformations of the ligand within the binding pocket, shedding light on the temporal evolution of this molecular interaction.

As previously described, procyanidin B2 presents one stable conformational state for approximately 220 ns, with an RMSD value ranging from 0.4 to 0.8 Å, and another state in the last 80 ns of simulation. As evident at 50 ns representation, NiV-G maintains this remarkably stable interaction with the ligand by adapting or narrowing its pocket’s surface. This adjustment effectively reduces the ligand’s degrees of freedom, ensuring a consistent and stable interaction over time. The interaction proportion plot of this procyanidin, highlights the central role of hydrophobic interactions in this biological event. As seen in **Figure** 9A, procyanidin B2 maintains nine hydrophobic interactions and eleven hydrogen bonds with the pocket. However, as the ligand transitions to its second preferred state, it loses two hydrophobic interactions with D219 and Y280 residues. As a consequence, the RMSD value of this ligand duplicates to 1.3 Å and the pocket cavity opens up a little reducing its constrained interaction with procyanidin B3.

In the case of procyanidin B3, as evident on the 3D representations (**Figure** 9B), the pocket cavity undergoes multiple dynamic adjustments as it interacts with the ligand. At the beginning of the simulation (**Figure** 9B) at 50 ns, procyanidin B3 is greatly constrained by the pocket cavity, exhibiting an RMSD value of 0.5 Å (**Figure** 4B). In this initial state, this ligand formed nine hydrophobic interactions and ten hydrogen bonds with NiV-G pocket. As the simulation progresses toward 60 ns, NiV-G pocket cavity starts to rearrange looking for a more favorable interaction with the ligand. Moreover, procyanidin B3 finds a stable conformational state that allows him to keep an RMSD value of almost 1.1 Å for almost 100 ns of simulation. This new binding pose observed between NiV-G pocket and the ligand is confirmed by the representation at 250 ns, and also, by the establishment of a new hydrogen bond (R248) with the pocket. As the simulation reaches its end, we observe a last binding pose with a NiV-G pocket more enclosed, covering the third preferred state of procyanidin B3 (**Figure** 9B at 290 ns).

This mechanism of ligand induced-fit adaptation of NiV-G pocket becomes even more evident with procyanidins B7 and C1. As indicated by earlier results, procyanidin B7 undergoes a first stable conformational state with RMSD values of 1Å. As it transitions to the second preferred state, it displays a more dynamic behavior until NiV-G pocket induced-fit conformational rearrange constrains its movement. This phenomenon is further confirmed by the tradeoff of interactions with NiV-G pocket residues. The first preferred state is maintained by eight hydrophobic interactions and eleven hydrogen bonds. Nevertheless, as NiV-G encloses around this ligand, some of the interactions are lost due to the less freedom degrees of procyanidin B7.

A further instance of this binding mechanism is evidenced with Procyanidin C1. However this adaptation process results in a more dynamic interaction that can explain its RMSD fluctuation in the last 100 ns (**Figure** 4D), but that ends with a more rigid and constrained conformation of NiV-G pocket (**Figure** 5D, **Figure** 9) at 250 ns. This loss of flexibility also leads to the disruption of multiple hydrogen bonds and hydrophobic interactions.

The remarkable stability of procyanidins within NiV-G pocket can be attributed to the robust interaction network mainly formed by hydrophobic interactions and hydrogen bonds. Notably, these results highlight the central role of D219, Y280, Y351, Y508, and K560 residues of NiV-G pocket for hydrophobic interaction with ligands, and also D302, G352, in addition to Y351 and Y508 for establishing the most hydrogen bonds during all the simulations.

Altogether, these results corroborate that the main protein-ligand interaction mechanism of NiV-G pocket with procyanidins B2, B3, B7, and C1 is an induced-fit conformational rearrange supported by hydrophobic and hydrogen bond interactions (**Figure** 8).

### 3.4. Binding free energy analysis

One of the key foundations of computational-guided drug discovery and design, is the estimation of binding affinities between ligand-receptor complexes. This value can be assessed through different approaches; however, the most important is the binding free energy 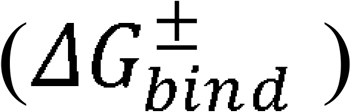 because it is directly related to the inhibition constant that can be measured experimentally [59].

Several methods have been developed to estimate this value, including the molecular docking approach used at the beginning of this study. Nevertheless, due to the limitations imposed by this method [60], we reconstructed the potential of mean force (PMF) from 7 independent replicates of SMD simulations with an external pulling force in the Z axis (see Methods 2.5), to calculate the 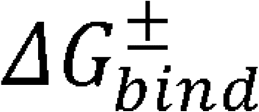 of each Procyanidin unbinding process.

This approach is based on two key concepts. First, SMD simulations allow us to introduce an external force that accelerates biological processes [61] such as the transition between bound and unbound states of procyanidins within NiV-G pocket. Secondly, following Jarzynski identity, we can estimate the free energy change between unbound and bound states based on ensemble average of the Boltzmann-weighted work performed during several non-equilibrium transformations from the initial to final states [37].

Our non-equilibrium 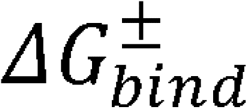 estimation revealed that Procyanidin B3 has the strongest binding affinity with NiV-G pocket with a value of 82.205 kcal·mol□^1^(**Figure** 10B), followed closely by Procyanidin B2 at 81.2474 kcal·mol□^1^(**Figure** 10A). Procyanidin B7 and Procyanidin C1 also exhibit substantial binding affinities although slightly lower with 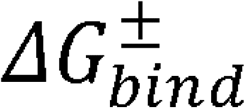 values of 74.5425 and 62.8547 kcal·mol□^1^respectively (**Figure** 10C-D).

**Figure 10.**
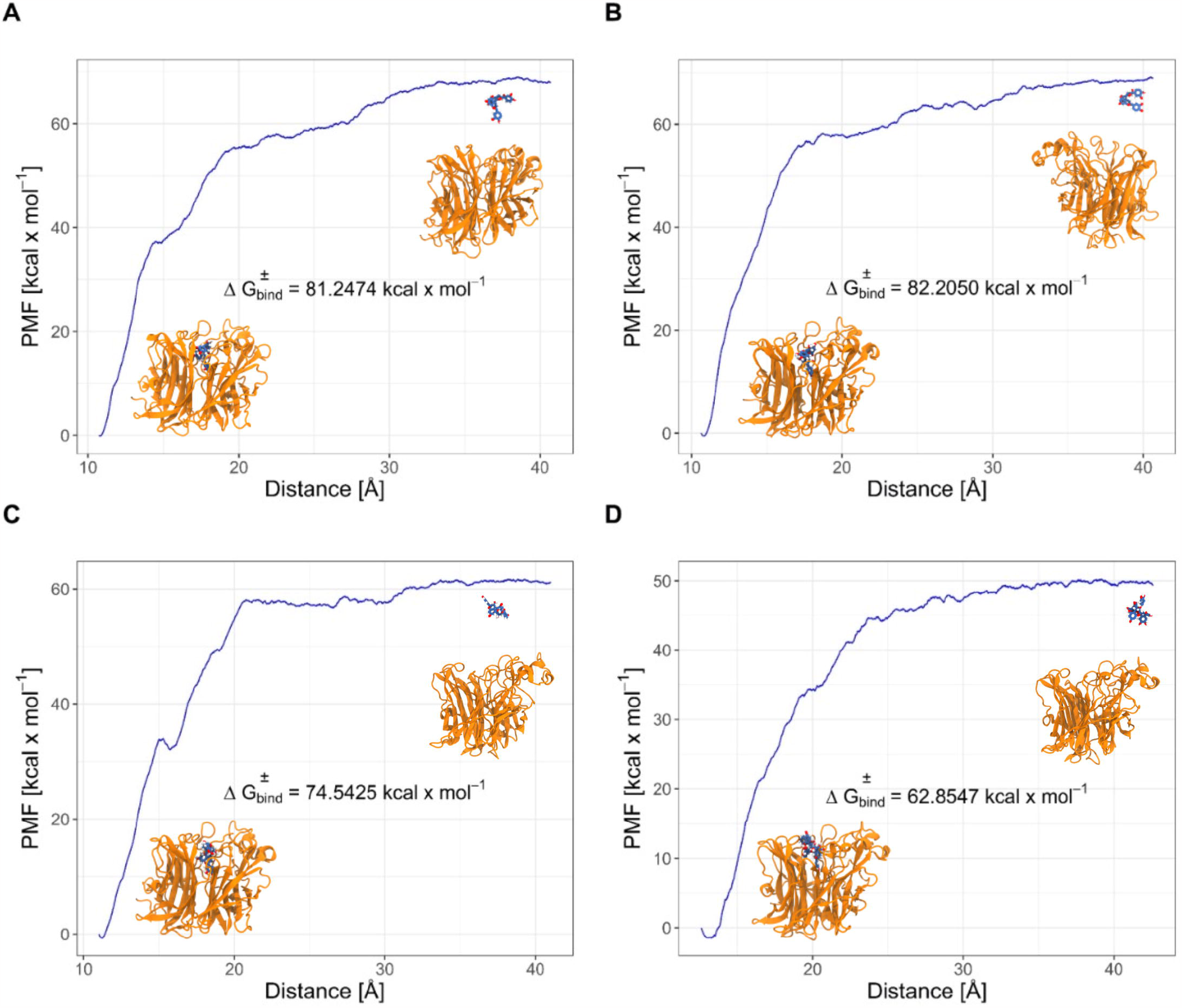
Non-equilibrium 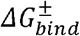 estimated for each procyanidin unbinding of NiV-G pocket. **A** PMF of Procyanidin B2 and NiV-G unbinding process. **B** PMF of Procyanidin B3 and NiV-G unbinding process. **C** PMF of Procyanidin B7 and NiV-G unbinding process. **D** PMF of Procyanidin C1 and NiV-G unbinding process. The 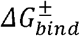 was calculated as described in methods.

Despite the difference on binding affinities order, these results are in accordance with our virtual screening results, which identify Procyanidin B3 as one the ligands with the highests affinity for NiV-G binding pocket. Overall, the procyanidins B2, B3, B7, and C1, represent promising lead compounds for future drug design against NiV-G.

## 4. Conclusions

This study employed computational biophysics to investigate Peruvian phytochemicals as potential inhibitors against the Nipah virus glycoprotein (NiV-G). Peruvian phytochemicals, particularly procyanidins B2, B3, and B7, demonstrated strong binding affinities and structural stability with NiV-G. These findings suggest the promise of these natural compounds as lead candidates for combating Nipah virus infections. The study’s insights emphasize the potential of natural compounds in drug discovery, offering hope for effective treatments against this global threat. Further research and experimental validation are essential to advance this work.

## 6. Declarations

### Data availability statement

All the data generated during the experiment are provided in the manuscript/supplementary material.

### Ethics approval statement

Not Applicable

### Patient consent statement

Not Applicable

### Permission to reproduce material from other sources

Not Applicable

### Clinical trial registration

Not Applicable

### Consent for participation/publication

Not Applicable

## Acknowledgements

The authors express their appreciation for the support provided the Laboratory of Clinical Genetics, Genomics, and Enzyme Research (LCGGER) within the Department of Genetic Engineering and Biotechnology, University of Chittagong, with special acknowledgment of the assistance offered by the International Foundation for Collaborative Research (IFCR). Also, we would like to acknowledge the support received from the Brazilian agencies CNPq, CAPES and FAPEMIG. Part of the results presented here were developed with the help of a CENAPAD-SP (Centro Nacional de Processamento de Alto Desempenho em São Paulo) grant UNICAMP/FINEP–MCT, CENAPAD– UFC (Centro Nacional de Processamento de Alto Desempenho, at Universidade Federal do Ceará), and Digital Research Alliance of Canada (via project bmh-491-09 belonging to Dr. Nike Dattani), for the computational support.

This study is dedicated to researchers worldwide who are committed to combatting Nipah virus infections. We extend our dedication to the dedicated researchers at the Institute of Epidemiology Disease Control and Research (IEDCR), under the leadership of the honorable Health and Family Welfare Minister of Bangladesh, Mr. Zahid Malek, MP. In addition, we dedicate this research to all the diligent professionals, students, and researchers striving to protect the world from the threat of infectious diseases.

## Author contributions

Conceptualization, GRP; Methodology, GRP, JRG; Software, GRP, JRG, EGV, and JPS; Validation, MCZ, KO, INA, HDW, and FSA; Formal analysis, GRP, JRG, EGV, and JPS; Investigation, GPR, JRG, EGV, and JPS; Resources, ND, IC, RBP, and ATM; Data curation, GRP, JRG, EGV, and JPS; Writing – original draft, GRP, and JPS; Writing – review and editing, GRP, JPS, RBP, and ATM; Visualization, EGV, and JPS; Supervision, GRP, and IC; Project administration, GRP, and IC; Funding acquisition, ND, IC, GRP, RBP and ATM. All the authors reviewed the manuscript and approved for submission.

